# Structure of the human taurine transporter TauT reveals substrate recognition and mechanisms of inhibition

**DOI:** 10.1101/2024.12.04.626611

**Authors:** Yishuo Lu, Dian Ding, Hongyi Chen, Peijun Jiang, Yuxin Yin

## Abstract

Taurine, a sulfur-containing amino acid, plays a crucial role in human health due to its antioxidant, anti-inflammatory, osmoregulatory properties ^1,2^. Taurine levels are primarily regulated by the sodium-/chloride-dependent taurine transporter (TauT) ^3^. Inhibitors of TauT have potential in the treatment of certain diseases, including neurological disorders and cancer ^4,5^. Here, we present five structures of TauT: in the substrate-free apo state, in complex with taurine and in complex with taurine-mimetic inhibitors β-alanine, GABA and guanidinoethyl sulfonate (GES), each with varying length of linear structure, in both inward-facing and occluded conformations. The taurine-, β-alanine- and GABA-bound hTauT structures, in the presence of NaCl, adopt an occluded conformation, with ligands binding in the central pocket. In the presence of KCl, GES-bound hTauT adopts an inward-facing conformation, with two molecules positioned along the substrate translocation pathway with one into the deep central cavity and the other precluded conformational change from inward-facing to occluded state. Combined with function analysis, our structures provided insights into the overall architecture, substrate coordination and inhibitor recognition mechanisms of TauT.

## Introduction

Taurine is a sulfur-containing non-protein amino acid that is found in high concentration in various excitatory and oxidative tissues, particularly in the brain, heart, skeletal muscles and retinal^1^. Taurine primarily resides in the intracellular fluid of these tissues, where its concentration can reach 10–15 mM, compared to approximately 50 μM in plasma^6^. Taurine participates in numerous physiological activities, including regulating cell volume, maintaining cell integrity^7,8^, forming bile salts^9^, reducing the risk of cardiovascular diseases^10^, promoting retinal differentiation^11^ and reducing neuronal apoptosis and inflammation^2^, through its membrane stabilizing, anti-oxidative and anti-inflammatory properties. Dysregulated taurine levels are associated with cardiomyopathy^12^, retinopathy^13^, neurological abnormalities^14^, weakened immune response^15^ and cancer^16,17^.

In most species, the biosynthesis of taurine occurs primarily in the liver and its biosynthetic capacity declines with aging ^3,18^. Thus, the high intracellular taurine levels are maintained by taurine intake through the action of the major taurine transporter, TauT (encoded by the *SLC6A6* gene)^3,19^. The expression level of TauT is high in the placenta and skeletal muscle, moderate in the heart, brain, lung and pancreas, and low in the liver ^5,20^. Loss-of-function mutations in TauT have been associated with dilated cardiomyopathy and retinal degeneration ^12,21,22^. On the other hand, overexpression of TauT has been observed in several types of cancers, such as gastric and colorectal cancer ^23,24^. High expression levels of TauT correlate with poor prognosis and advanced tumor stages, suggesting that TauT may be a potential target for anticancer therapies ^5,25^.

TauT is a chloride- and sodium-dependent transporter that belongs to the solute carrier 6 (SLC6) family. This family can be divided into four subgroups: GABA (γ-aminobutyric acid), monoamine, neurotransmitter amino acid, and nutrient amino acid transporters. TauT belongs to the first subgroup based on the similarity of its substrates (Extended Data Fig. 1a) ^5^. Progress in structural studies of SLC6 transporters has revealed competitive and allosteric mechanism of their inhibitors ^26–32^. Inhibition of GAT1 (SLC6A1) dependent GABA clearance by tiagabine in the synaptic cleft is an established strategy for treating epilepsy and structural studies elucidate the mode of action of these inhibitors and provides blueprints for the design of neuromodulators ^33,34^. RGX-202, an inhibitor of the creatine transporter (SLC6A8), is under clinical trials for colon cancer treatment ^5,35^. However, there is limited information about TauT inhibitors. Although several substrate-mimetics inhibitors of TauT, with simple or more complex structure, have been reported, high resolution structure of TauT may largely contributed to the development of highly selective and potent TauT inhibitors with anti-neoplastic property ^36,37^.

Here we report five structures of TauT in the apo state, in complex with its substrate taurine and in complex with its inhibitors with different length of linear structure in the inward-facing and occluded conformations. Combine with function analysis, these structures reveal the overall architecture, substrate binding and inhibitor recognition mechanism of TauT.

## Results

### Overall architecture of human TauT

Xenopus laevis oocytes injected with human TauT cRNA exhibited time-dependent accumulation of [^3^H]-taurine (Extended Data Fig. 1b) with an apparent IC_50_ of 2.63 ±1.04 µM (Extended Data Fig. 1c). Full-length wild-type hTauT, fused with a C-terminal affinity tag, was expressed in HEK293F cells^38^. Recombinant hTauT protein was extracted from membranes using lauryl maltose neopentyl glycol (LMNG) and cholesteryl hemisuccinate (CHS). The affinity tag was removed, and hTauT was purified using size exclusion chromatograph (SEC) (Extended Data Fig. 1d). Protein from the peak fractions was used to prepare cryo-EM samples, leading to the determination of hTauT structures in the apo state (hTauT_APO_), in complex with substrate taurine (hTauT_TAU_), and in complex with inhibitors GABA (hTauT_GABA_), β-alanine (hTauT_βA_) and guanidinoethyl sulfonate (GES) (hTauT_GES_). The resolutions of these structures range from 2.9 Å and to 3.3 Å (Fig. 1, Extended Data Fig. 2-4 and Extended Data Table 1). The structures of hTauT in the apo state, in complex with taurine, β-alanine and GABA, were prepared in the presence of 150 mM NaCl, while the GES-bound structure was prepared in the presence of 120 mM KCl. These five structures exhibit inward-open and occluded conformations.

**Figure 1.**
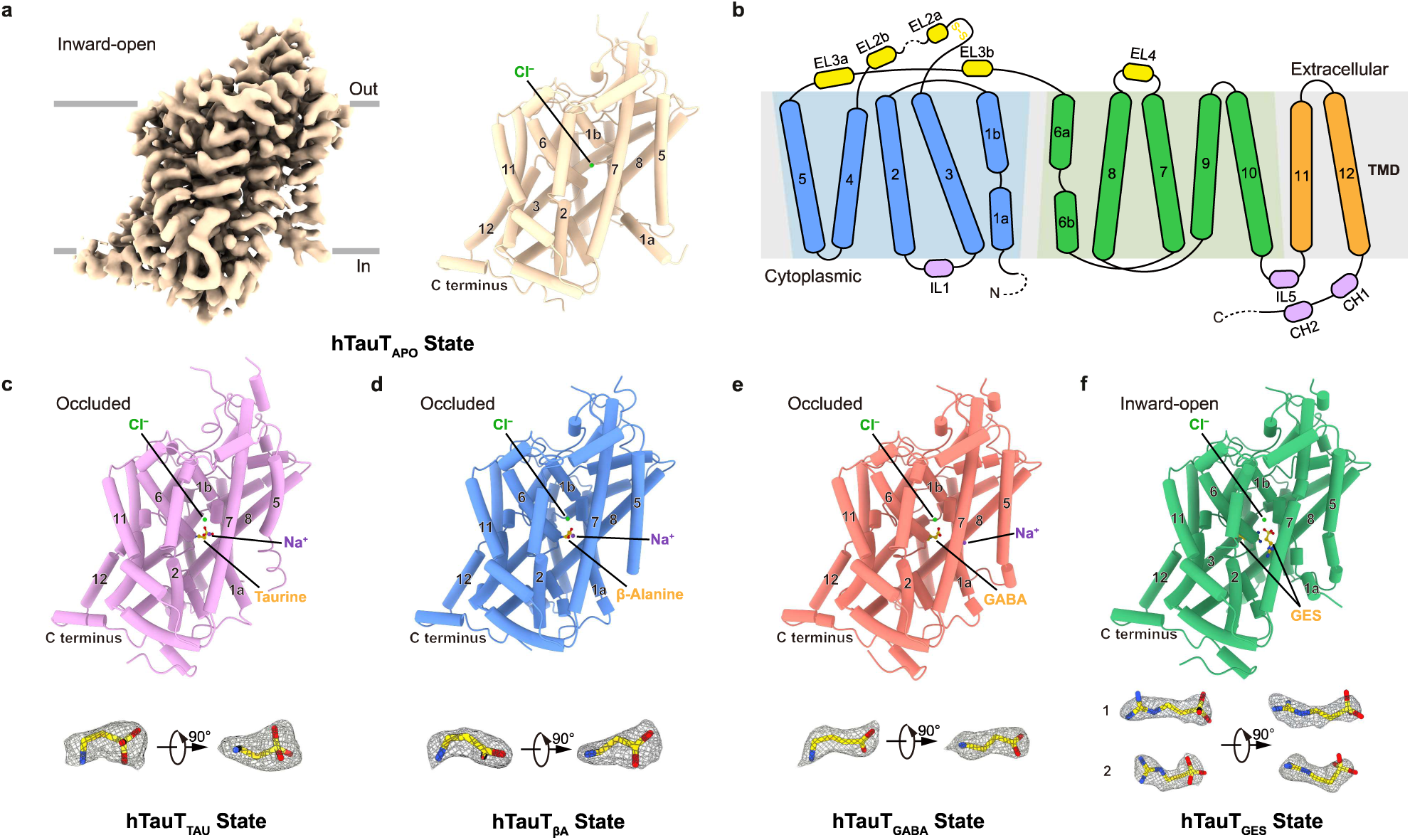
Cryo-EM structures of human TauT. **a,** Cryo-EM density and the overall structure of human TauT (hTauT) in the apo state. The Cl^−^ ion is represented as green dot. **b,** Topology of the secondary structure of hTauT. TM1-TM5 are colored in blue and TM6-10 with inverted symmetry are colored in green. Extracellular and intracellular helices are colored in yellow and pink, respectively. Dashed lines indicate the disordered regions in the cryo-EM structure. **c-e,** Structures of hTauT in complex with taurine (c), β-Alanine (d) and GABA (e) in the presence of NaCl. Densities of ligands are shown as mesh. **f,** Structures of hTauT in complex with guanidinoethyl sulfonate (GES) in the presence of KCl. Densities of two GES molecules are shown as mesh.

The structure of hTauT_APO_ features a canonical LeuT fold with pseudo-twofold symmetry organization of TMs 1–5 relative to TMs 6–10 and the transmembrane domain displays an inward-opening (Fig. 1a and 1b) conformation. The central substrate-binding site is accessible from the cytoplasmic side (Extended Data Fig. 5a). Density of the N-terminal sequence and TM1a (residue 1-57) was less clear in the inward-opening state due to flexibility (Extended Data Fig. 4b). Intracellular loops (ILs) and extracellular loops (ELs) except for EL2 (residue 180-188), connecting the TMs were unambiguously modelled into the density. The closed extracellular gate is formed by interactions between residues from TM1b, TM3, TM6a and TM10 (Extended Data Fig. 5b). Density of Cl^−^ was observed at the conserved chloride-binding site (Extended Data Fig. 5c). Ligand-bound hTauT structures in the presence of NaCl all adopt occluded states, with one molecule binding in the central binding pocket formed by the unwound regions of TM1 and TM6, together with adjacent residues in TM3 and TM8 (Fig. 1c-1e). The GES-bound structure in the presence of KCl adopts an inward-facing conformation, with two molecules positioned within the central cavity (Fig. 1f).

### Taurine and ion binding pockets

Human TauT (hTauT) exhibited taurine transporter activity with a *K*_m_ of 7.62 ± 1.96[µM (Fig. 2a). The structure of hTauT in complex with taurine adopts an occluded conformation at an overall resolution of 3.3 Å (Fig. 2b and 2c, Extended Data Fig. 3b). The superposition of taurine-bound hTauT with hTauT_APO_ shows overall root mean-square deviation (r.m.s.d.) values of 1.86 Å, with TM1a undergoing an obvious conformational change that occludes the intracellular permeation pathway (Fig. 2b and 2c). TM1, TM3, TM6 and TM8 enclosed the central substrate-binding pocket S1 (Fig. 2d). Density in the pocket was identified as taurine molecule and additional densities of Na^+^ and Cl^−^ ions were observed in the conserved Na1 and chloride binding site ^39^ (Fig. 2d). According to the three subsite (A, B and C) representation of the central binding pocket, taurine occupies subsite A with its sulfate group and a water molecule was observed in subsite C (Fig. 2d). One oxygen atom of the sulfate group of taurine coordinates Na^+^, one oxygen atom forms hydrogen bond with nitrogen atom of Gly62 and the other oxygen atom forms hydrogen bond with hydroxyl group oxygen of Ser402 and nitrogen atom of Gly60 (Fig. 2e). The amino group of taurine forms a hydrogen bond with carbonyl group of F300, while the carbon linker is stabilized by the side chain of Tyr138 and Phe300 (Fig. 2e). Na^+^ at Na1 site is coordinated by the side chains of Asn63, Asn333, Ser301 and carbonyl group of Phe58 and Ser301 (Fig. 2f, Extended Data Fig. 6a). The Cl^−^ is chelated by the side chains of Tyr83, Gln297, Ser337 and Ser301 (Fig. 2g, Extended Data Fig. 6d). No density corresponding to Na^+^ ion was observed at the conserved Na2 site, however a taurine like density was identified near the Na2 site, surrounded by TM1 (Val59), TM5 (Ala250 and Phe254) and TM8 (Leu398 and Ser402) (Extended Data Fig. 6g).

**Figure 2.**
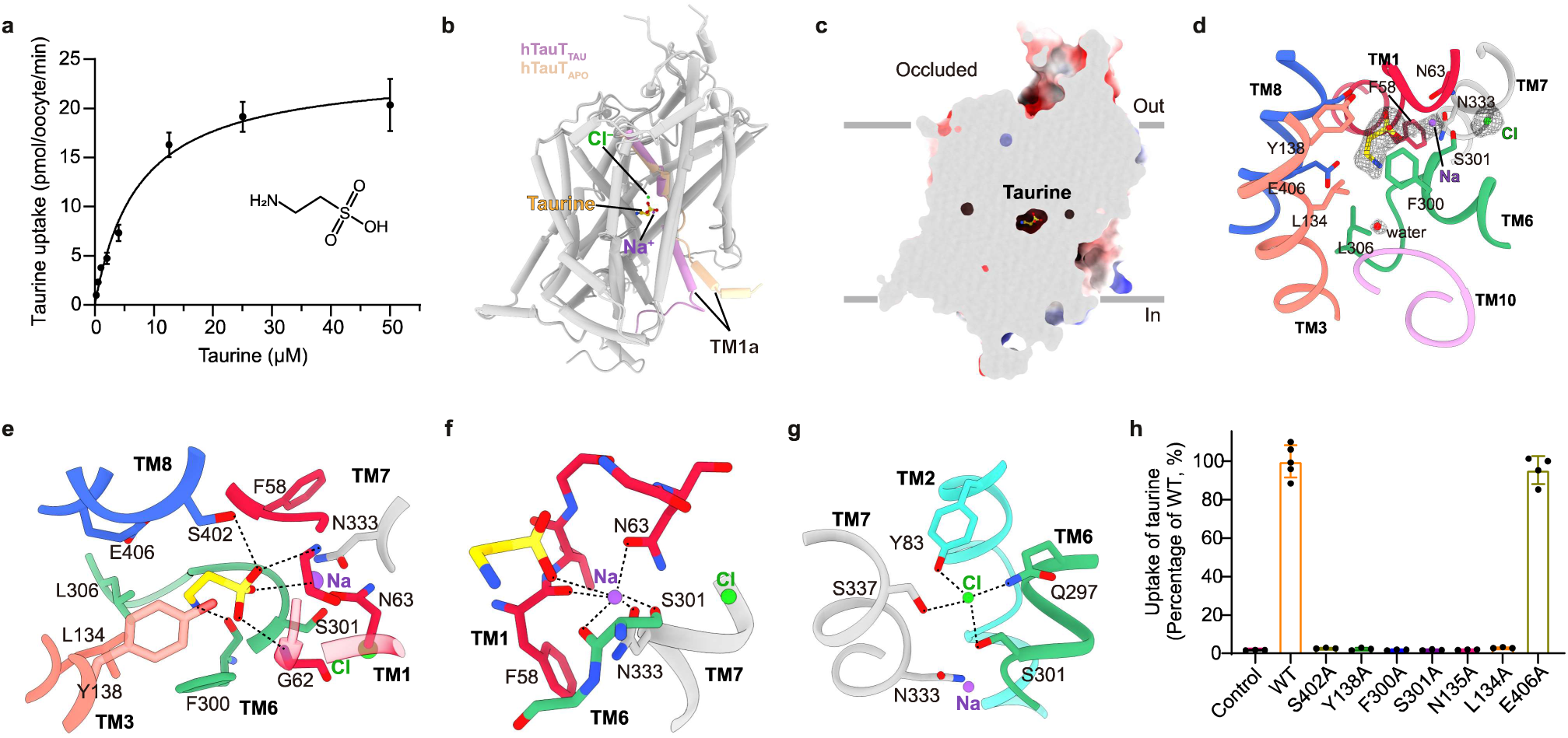
Taurine and ion binding site of hTauT. **a,** The transport kinetic of taurine uptake of hTauT. The Michaelis constant (Km) value and Vmax for [^3^H]taurine uptake is 7.62 ± 1.96[μM and 24.28 ± 2.04 pmol/oocyte/min. Data are shown as mean ± SD (n=3 biologically independent experiments). **b,** Structure of taurine-bound hTauT superimposes with hTauT_APO_. TM1 of hTauT_TAU_and hTauT_APO_ are colored in pink and light peach, repectively. Taurine is shown as sticks. Na^+^ and Cl^−^ ions are shown as dot. **c,** Cut-open view of electrostatic surface of hTauT_TAU_ with taurine binds in the central pocket. **d,** Model of the taurine-binding pocket. Cryo-EM densities of taurine, Na^+^, Cl^−^ and water are shown as mesh. Taurine is shown as sticks. Na^+^, Cl^−^ and water are shown as dot. **e,** Close-up views of taurine at substrate binding pocket. Taurine and surrounding residues are shown as stick and Cl^−^ and Na^+^ ions are shown as dot. The coordination of taurine is indicated by dashed lines. **f,** Close-up views of Na^+^ at Na1 site. Key residues and taurine coordinate Na ion are shown as stick and Cl^−^ and Na^+^ ions are shown as dot. The coordination of Na is indicated by dashed lines. **g,** Close-up views of the chloride binding site. Key residues coordinate Cl^−^ ion are shown as stick and Cl^−^ and Na^+^ ions are shown as dot. The coordination of Cl is indicated by dashed lines. **h,** Effects of alanine substitution of key residues in the taurine-bound pocket on transport activity of hTauT. Data are expressed relative to wild-type hTauT and the bars represent mean ± SD (n≥3 biologically independent experiments).

To validate the function of residues surrounded taurine and ions, mutagenesis experiments were performed by measuring the uptake of radiolabeled taurine of TauT mutants. Substitutions of S402A, Y138A and F300A which coordinate taurine leads to complete loss of the transport activity (Fig. 2h). Substitution of Ser301, which coordinates Cl^−^ and Na^+^ at Na1 site, to alanine also abolished the transport of taurine (Fig. 2h). E406A mutation at subsite B do not affect the transport activity of TauT, however, substitution of L134A and N135A completely abolish taurine transportation (Fig. 2h).

### Binding mode of **β**-alanine

Previous results have revealed that the best inhibitor for TauT is linear substrate analogues especially β-alanine which have a three-carbon linker and a amine group identical to taurine ^5^. β-alanine shows inhibitory activity with IC_50_ values of 31.65 ± 10.76 µM (Fig. 3a), which is similar to the value measured using HEK293 cells ^36^. The change of acidic moiety to carboxy group decreased the inhibitory effect 12-fold compared with sulfate group (Extended Data Fig. 1c). The structure of hTauT_βA_ adopts occluded conformation which is almost identical to hTauT_TAU_ with the overall r.m.s.d. values of 1.16 Å (Fig. 3b-3c and 3e-3f). β-alanine binds at the same site as taurine at subsite A and densities of water, Na^+^ and Cl^−^ were observed at conserved subsite B, Na1 and chloride-binding site, respectively (Fig. 3d, Extended Data Fig. 6b and 6e). One oxygen atom of the carboxy group forms hydrogen bond with carbonyl group of Ser301 and the other oxygen coordinates Na^+^ and form hydrogen bond with nitrogen atom of Gly62 (Fig. 3d). The amino group of β-alanine also forms hydrogen bond with carbonyl group of F300 and the carbon linker is stabilized by the side chain of Tyr138 and Phe300 (Fig. 3d). At conserved Na2 site, a β-alanine like density was observed similar to the density of hTauT_TAU_ (Extended Data Fig. 6h).

**Figure 3.**
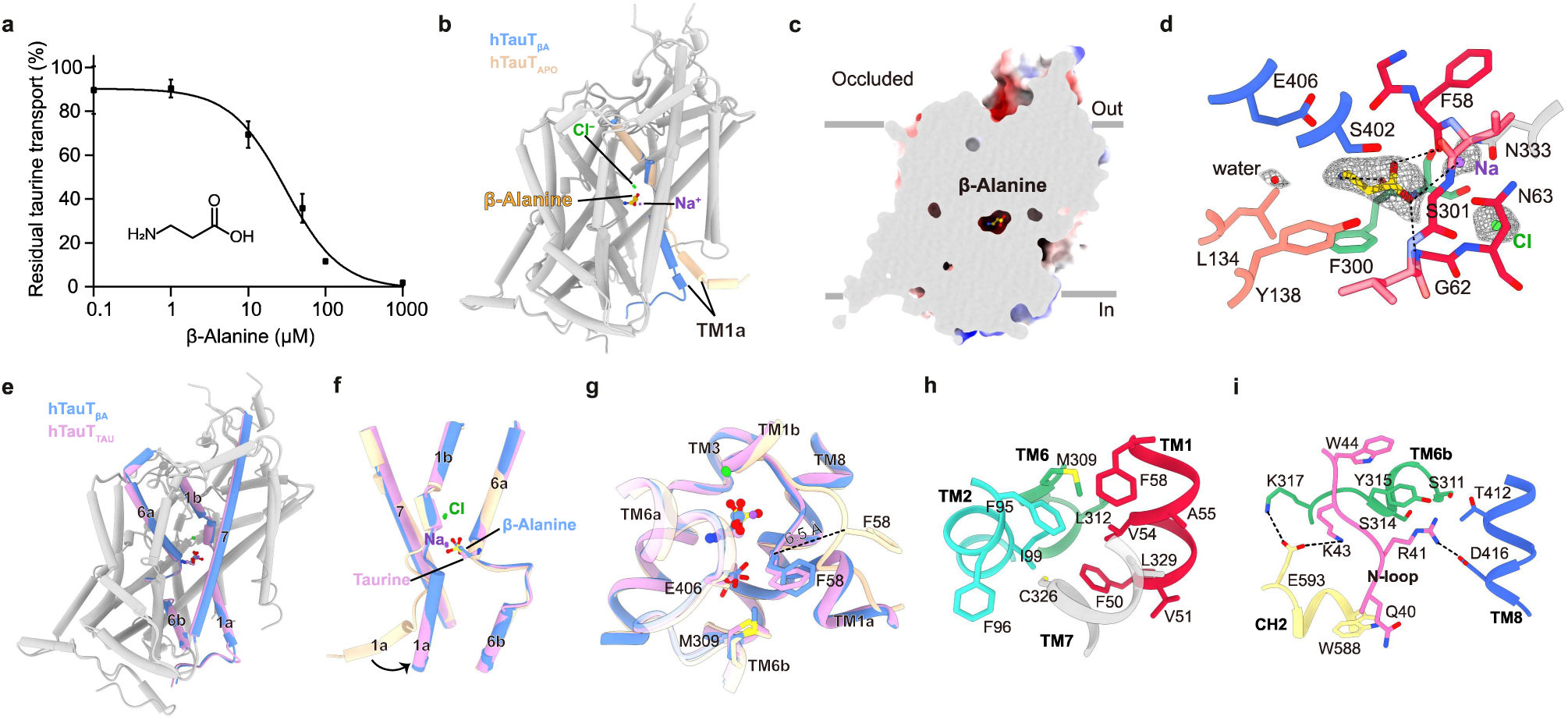
Binding mode of β-alanine and structural comparison with inward-facing conformation. **a,** Effect of β-alanine on taurine uptake by hTaut in oocytes. The IC_50_ value for β-alanine is 31.65 ± 10.76[µM. Data are shown as mean ± SD (n=3 biologically independent experiments). **b,** Structure of β-alanine-bound hTauT superimposes with hTauT_APO_. TM1 of hTauT_βA_ and hTauT_APO_ are colored in blue and light peach, respectively. β-alanine is shown as sticks. Na^+^ and Cl^−^ ions are shown as dot. **c,** Cut-open view of electrostatic surface of hTauT_βA_ with β-alanine binds in the central pocket. **d,** Close-up view of the β-alanine-binding site. Densities of β-alanine, Na^+^, Cl^−^ and water are shown as mesh. β-alanine is shown as sticks. Na^+^, Cl^−^ and water are shown as dot. The coordination of β-alanine is indicated by dashed lines. **e,** Superposition of hTauT_βA_ with hTauT_TAU_. TM1 and TM6 of hTauT_βA_ and hTauT_TAU_ are colored in blue and pink, respectively. β-alanine and taurine are shown as sticks. **f,** Structural comparison of hTauT_APO_ in the inward-facing conformation with hTauT_βA_ and hTauT_TAU_ in the occluded conformation. The arrow indicates the movement of TM1a from inward-facing to occluded conformation. The black lines indicate the same binding mode of taurine and β-alanine. Na^+^ and Cl^−^ ions are shown as dot. **g,** Detailed conformational change of gating residues. Residues preclude ligand release are shown as sticks. Distance between the Cα of F58 of hTauT_TAU_ and hTauT_APO_ is indicated by dashed line. **h-i**, Hydrophobic interactions (h) and salt bridges (i) formed the intracellular gate. Salt bridges are indicated by dashed lines.

The major differences between the substrate-free and substrate-bound hTauT is TM1a and N-terminal loop (Fig. 3f). The conformational change of TM1a of hTauT_APO_ swings F58 away from the pocket, makes the central pocket accessible from the cytoplasmic side (Fig. 3f-3g). This movement disrupts the Na1 binding site and leaves the chloride binding site unaffected (Extended Data Fig. 6j and 6l). Hydrophobic interaction between residues of TM1 (Phe50, Val51, Val54, Ala55 and Phe58), TM2 (Phe95, Phe96 and Ile99) and TM6 (Met309 and L312) and TM7 (L329) stabilized the occluded conformation (Fig. 3h). The clear N-terminal and C-terminal densities of hTauT_TAU_ enabled us to identify the salt bridges formed at the occluded conformation (Extended Data Fig. 4a). Interaction between side chain of Arg41-Asp416 and Lys317-Glu593-Lys43 further stabilized the occluded conformation (Fig. 3i).

### Coordination of GABA

GABA, the linear inhibitor TauT, is different from β-alanine with a one atom longer carbon linker (Fig. 4a). Inhibitory effect of GABA has been reported with IC_50_ values of 1014 µM and the interaction of GABA-mimetics with TauT has been investigate^37^. The IC_50_ value of GABA for inhibiting hTauT is 66.42 ± 53.56 µM, around 15 times more potent than previous reported (Fig. 4a). Comparing with β-alanine, longer carbon linker does not significantly decrease the inhibitory effect of GABA (Fig. 3a and 4a). GABA-bound hTauT is in the occluded conformation with GABA molecule binds at the central cavity (Fig. 4b and 4c). GABA bind at subsite A with its carboxy group and its amino group extends toward subsite B, and densities of water molecules were observed at subsite B and C (Fig. 4d). Densities of Na^+^ and Cl^−^ were observed at conserved Na2 site and chloride-binding site (Fig. 4d, Extended Data Fig. 6f and 6i). No clear Na^+^ density and ligand-like density were observed at Na1 site and near Na2 site as observed in maps of hTauT_TAU_ and hTauT_βA_ (Extended Data Fig. 6c and 6g-6i).

**Figure 4.**
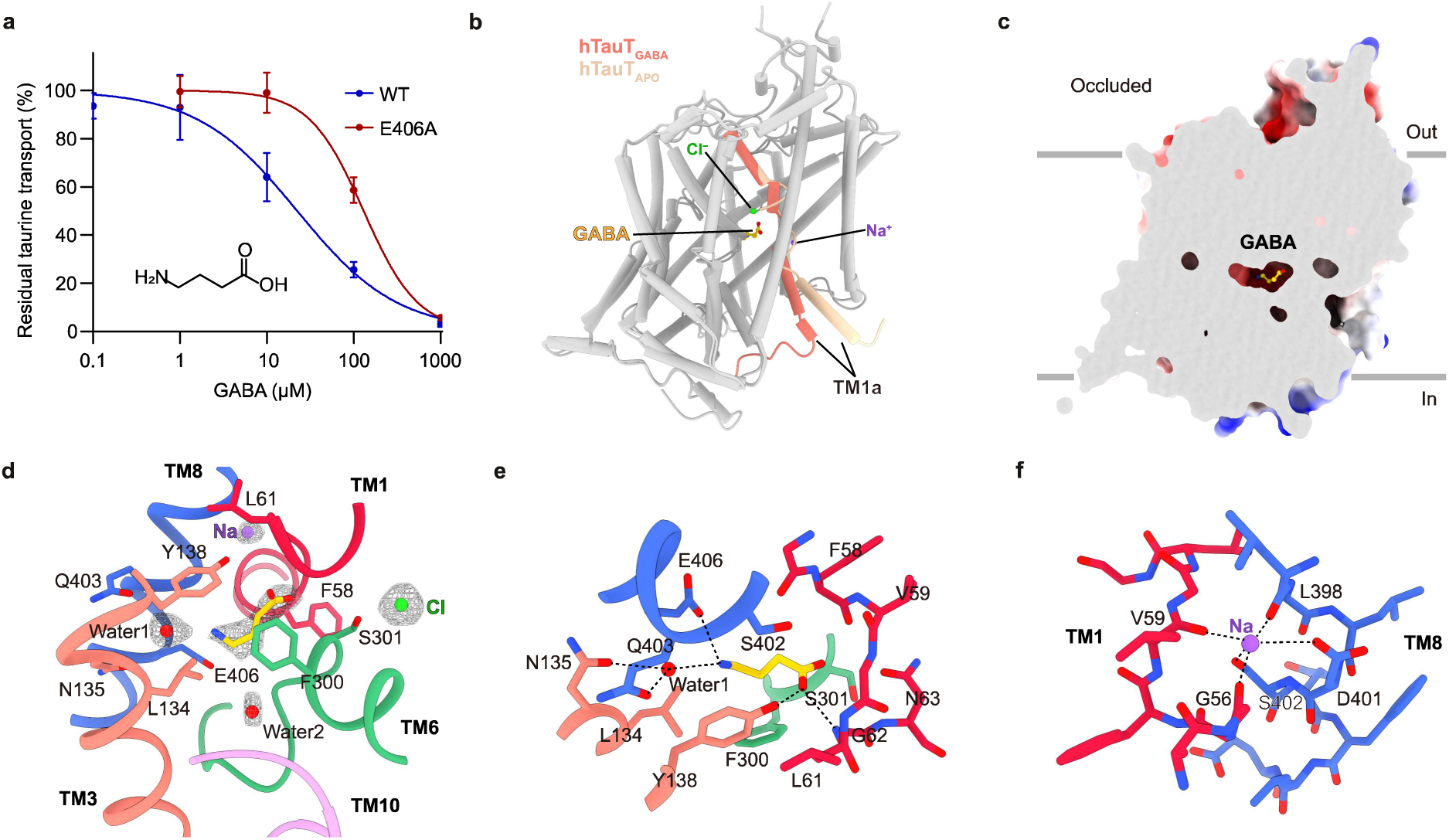
Recognition of GABA in the central pocket. **a,** Effect of GABA on taurine uptake by wild-type and E406A mutant hTauT in oocytes. The IC_50_ value for GABA of wild-type and E406A mutant hTauT are 66.42 ± 53.56[µM and 134.28 ± 25.92[µM. Data are shown as mean ± SD (n=3 biologically independent experiments). **b,** Structure of GABA-bound hTauT superimposes with hTauT_APO_. TM1 of hTauT_GABA_ and hTauT_APO_ are colored in orange and light peach, respectively. GABA is shown as sticks. Na^+^ and Cl^−^ ions are shown as dot. **c,** Cut-open view of electrostatic surface of hTauT_GABA_ with GABA binds in the central pocket. **d,** Model of the GABA-binding pocket. Cryo-EM densities of GABA, Na^+^, Cl^−^ and water are shown as mesh. GABA is shown as sticks. Na^+^, Cl^−^ and water are shown as dot. **e,** Close-up views of GABA at substrate binding pocket. GABA and surrounding residues are shown as stick and Cl^−^ and Na^+^ ions are shown as dot. The coordination of GABA is indicated by dashed lines. **f,** Close-up views of Na^+^ at Na2 site. Key residues coordinate Na^+^ ion are shown as stick and Na^+^ ion is shown as dot. The coordination of Na is indicated by dashed lines.

The carboxy group of GABA forms a hydrogen bond with nitrogen of Gly62 and the hydroxyl group of Tyr138 (Fig. 4e). The amino group of GABA forms a hydrogen bond with the side chain of Glu406 and a water molecule (Fig. 4e). Na^+^ at the Na2 site is coordinated by the side chains of Asp401 and Ser402, the carbonyl oxygens of Gly56, Val59 and Leu398 (Fig. 4f). Superposition of hTauT_APO_ with hTauT_GABA_ shows the carbonyl group of Gly56 swings away from the binding pocket during occluded to inward-facing conformational change (Extended Data Fig. 6k).

Comparing with the central binding pocket of GAT1 shows that residues consist of subsite A are identical (Extended Data Fig. 1a and 7a). Substitution of Y60G and G279A from GAT1 to TauT may reduce the binding affinity of GABA which is consistent with previously reported functional experiments ^33^. Substitution of Glu406 to alanine, which does not affect the binding of taurine, reduced the inhibitory effect of GABA (Fig. 4a). The GABA coordination function of negatively charged glutamate is supported by the fact that Tyr60 (corresponding to Gly57 near Glu406) is substituted to glutamate in GAT2 and GAT3. Inhibitory effect of nipecotic acid, a GABA-mimetic, was reported with IC_50_ values of 2.02[mM ^37^. Comparison of hTauT_GABA_ with occluded nipecotic acid-bound GAT1 demonstrates that nipecotic acid may be coordinated by hTauT in a similar mode (Extended Data Fig. 7b).

### Inhibition of guanidinoethyl sulfonate

Guanidinoethyl sulfonate (GES), a substrate-mimetics of TauT, acts as a competitive inhibitor of taurine transport and has been reported effectively reduced taurine level in brain and plasma ^40,41^. GES shows inhibitory activity with IC_50_ values of 5.27 ± 1.46[µM, similar to the effect of taurine (Fig. 5a). GES-bound hTauT sample was prepare in the presence of KCl and the structure adopts an inward-facing conformation with the central pocket accessible from the cytoplasmic side (Fig. 5b and 5c). The binding mode of GES is different from other substrate analogues with two molecules binding at the central pocket (Fig. 5b and 5c).

**Figure 5.**
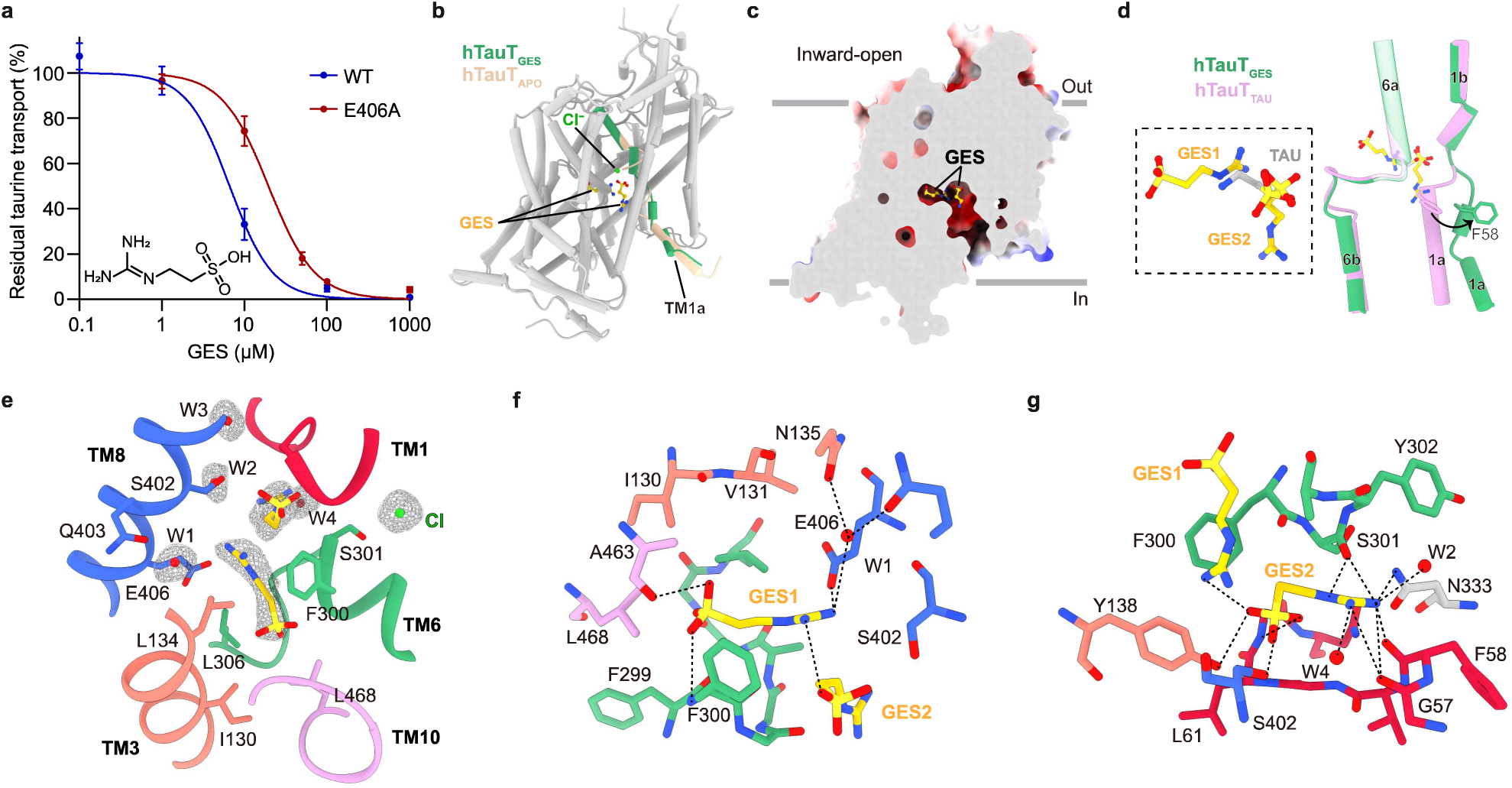
Structural basis for inhibition of GES. **a,** Effect of GES on taurine uptake by wild-type and E406A mutant hTauT in oocytes. The IC_50_ value for GES of wild-type and E406A mutant hTauT are 5.27 ± 1.46[µM and 19.68 ± 2.42[µM. Data are shown as mean ± SD (n=3 biologically independent experiments). **b,** Structure of GES-bound hTauT superimposes with hTauT_APO_. TM1 of hTauT_GES_and hTauT_APO_ are colored in green and light peach, repectively. GES is shown as sticks. Cl^−^ ion is shown as dot. **c,** Cut-open view of electrostatic surface of hTauT_GES_ with two GES molecules bind in the central pocket. **d,** Comparison of ligand binding pocket of hTauT_GES_ and hTauT_TAU_. **e,** Model of the GES-binding pocket. Cryo-EM densities of GES, Cl^−^ and water are shown as mesh. GES is shown as sticks. Cl^−^ and water are shown as dot. **f,** Close-up views of GES1 at substrate binding pocket. GES1, GES2 and surrounding residues are shown as stick. Water molecule is shown as dot. Hydrogen bonds form between residues, water and GES are indicated by dashed lines. **g,** Close-up views of GES2 at substrate binding pocket. GES1, GES2 and surrounding residues are shown as stick. Water molecules are shown as dot. Hydrogen bonds form between residues, water and GES are indicated by dashed lines.

Superposition of hTauT_GES_ with hTauT_TAU_ shows that the conformational change of TM1a leads to the opening of the binding pocket of the second GES molecule (Fig. 5d). Guanidine group of GES1 occupies at the same site as the amino group of taurine, while the sulfate group of GES2 overlaps with the sulfate group of taurine (Fig. 5d). Several densities of water molecules were observed surrounded GES molecules (Fig. 5e). Density of GES1 located within the central pocket with sulfate group binds at subsite C and guanidine group coordinate by water molecule at subsite B (Fig. 5e). GES2 is parallel to the substrate translocation pathway with its sulfate group occupying subsite A and guanidine group facing the cytoplasmic side (Fig. 5e). Sulfate group of GES1 forms hydrogen bonds with the back bone of Phe300 and Ala463, guanidine group interact with Glu 406, sulfate group of GES2 and water molecule coordinated by the side chain of Asn135 and Gln403, and carbon linker is coordinated by Phe300, Ala303 and Leu306 (Fig. 5f). Similar to the binding mode of acidic group of taurine, sulfate group of GES2 occupies subsite A. Oxygens of GES2 form hydrogen bonds with GES1, Gly62, Tyr138 and Ser402 (Fig. 5g). Guanidine group is coordinated by hydrogen bond network formed between Ser301, Asn333, Gly57, Phe58 and water molecules (Fig. 5g). The substitution of E406A reduced the inhibitory effect of GES, similar to GABA, demonstrates the function of Glu406 in the coordination of ligand with longer linker between acidic group and basic group (Fig. 5a). Comparison with tiagabine-bound GAT1 in the inward-facing conformation shows that sulfate group of GES2 occupies the same site as carboxy group of tiagabine and the movement of TM1a opens the binding pocket of the groups on the other end of these inhibitors (Extended Data Fig. 7c).

## Discussion

Here we presented the structures of human TauT in the apo state, in complex with taurine and in complex with substrate-mimetic inhibitors with different acidic groups, basic groups and lengths of carbon linkers. Comparing to the inward-facing conformation of substrate-free hTauT_APO_, the structure of hTauT_TAU_ reveals the central binding pocket and the coordination mode of Na^+^ at the Na1 site and Cl^−^ at conserved chloride-binding site, similar to GABA-bound GAT1^33^. Residues in subsites B and C, which do not directly coordinate taurine, may participate in shaping the central cavity and the coordination of water molecules. Water molecules in the cavity help coordinate the substrate by forming hydrogen bonds and acting as proton donors, facilitating the hydrolysis of the basic tail and the formation of cation-π interaction between taurine and Tyr138 and Phe300. The presence of Na^+^ and the substrate stabilizes hTauT in the occluded state, as the replacement of NaCl with KCl leads to the inward-facing conformation of hTauT_GES_. Taurine- and β-alanine-like densities observed near the conserved Na2 site probably stabilize the occluded conformation, mimicking the function of Na^+^ at Na2 site and preventing the release of substrates to the cytoplasmic side, as high intracellular taurine levels reduce transport activity of TauT ^3,42^.

The elevated expression level of the TauT transporter is associated with poor prognosis and advanced tumor stage in various cancers, suggesting the importance of taurine in cancer development ^5,23–25^. Recent study reported the pro-tumoral role of taurine and indicated that its anticancer effects in earlier studies might have been due to the restoration of antitumor immunity, which is impaired by the exhaustion of environmental taurine depleted by tumor cells ^43^. Inhibition of taurine transportation could be a promising strategy in anticancer therapy. The structures of hTauT_βA_, hTauT_GABA_ and hTauT_GES_ reveals the binding modes of these linear inhibitors. All of these molecules occupy subsite A with their acidic heads. Taurine includes an ethane backbone with one amino group attached to one carbon and one sulfate group attached to the other carbon. The length of β-alanine is similar to taurine, and they bind at the same site in the central pocket of hTauT. Functional studies indicate that the sulfate group probably enhance the affinity of ligands by donating protons, which facilitates the coordination of Na^+^ at Na1 site and interact with more residues in subsite A. In comparison, GABA, with a four-carbon backbone, has a similar inhibitory effect to β-alanine. The amino tail of GABA is coordinated by Tyr138 and Phe300 through cation-π interaction and Glu406 facilitates the binding of zwitterion with longer carbon linkers. GES, a more complex structure, shows potent inhibition of hTauT and may have unexplored antitumor properties. Base on the binding mode of GES, additional groups attached to the amino group of taurine could be accommodated in subsite C, providing a blueprint for developing more effective inhibitors. Our findings provide a detailed structural basis for understanding TauT function and its inhibition, offering valuable insights for future therapeutic development.

## Methods

### Cell lines

FreeStyle 293 F (Thermo Fisher Scientific) suspension cells were cultured in 293 Expression Medium (Gibco) supplemented with 1% FBS at 37 °C, in an atmosphere of 6% CO_2_ and 60% humidity. Sf9 insect cells (Thermo Fisher Scientific) were cultured in Sf-900 III SFM medium (Thermo Fisher Scientific) at 27 °C.

### Oocyte expression systems and flux experiments

*Xenopus laevis* oocytes were isolated and maintained as previously described^44^. Briefly, ovarian fragments were isolated from the animal and digested with collagenase (2mg/ml, type IA, Sigma-Aldrich) for 2 h in OR2 buffer containing 82.5 mM NaCl, 2 mM KCl, 1 mM MgCl_2_ and 5 mM Hepes (pH 7.4). Oocytes were then washed in ND96 buffer containing 96 mM NaCl, 2 mM KCl, 1.8 mM CaCl_2_, 1 mM MgCl_2_ and 5 mM HEPES (pH 7.8). Stage VI oocytes were transferred to fresh OCM containing 60% Leibovitz L15, 100 μg/ml gentamycin and 10 mM Hepes (pH 7.8) and were maintained at 18 - 20 °C prior to injection. cDNAs of wild-type and mutant hTauT were transferred into the pKSM vector. cRNA was obtained using the T3 mMessage mMachine Kit (Thermo Fisher Scientific) and 25 ng injected into selected oocytes. Oocytes were incubated at 20 °C for 40-48 h in ND-96 solution with 100 μg/ml gentamycin to allow expression of TauT. Uptake was measured in buffer containing 100 mM NaCl, 2 mM KCl, 1 mM CaCl_2_, 1 mM MgCl_2_ and 1 mM HEPES (pH 7.4) and medium containing the desired amino acid (Thermo Fisher Scientific) concentration and 10 μCi/ml [^3^H]taurine (PerkinElmer). For the assessment of time-dependent accumulation of taurine, oocytes were incubated in buffer containing 4 μM taurine (10 μCi/ml). For assessment of mutant TauT activity relative to wild-type as well as the inhibitory effects of inhibitors, oocytes were incubated in buffer containing 0.25 μM taurine (10 μCi/ml). Oocytes were solubilized by adding 1% Triton followed by liquid scintillation fluid for to quantitate radioactivity. Activity of TauT were calculated by subtracting the uptake of the non-injected group from that of the cRNA-injected groups.

### Expression and purification of TauT

cDNA encoding human wild-type TauT was cloned into the BacMam expression vector with C-terminal GFP-His-Strep tags^45^. The baculoviruses were produced using the Bac-to-Bac system and amplified in Sf9 cells. For protein expression, HEK293F cells cultured in Freestyle 293 medium at a density of 2.5[×[10^6^[ml^−1^ were infected with 10% volume of virus. Sodium butyrate (10 mM) was added to the culture 8 h post-infection and cells were harvested 60 h post-infection. Cells were collected by centrifugation at 3,724g for 10 min and washed with 20 mM HEPES (pH 7.4) containing 150 mM NaCl.

Cell pellets were solubilized in buffer containing 150 mM NaCl, 20 mM Hepes at pH 7.4 (HBS), 1% (w/v) lauryl maltose neopentyl glycol (LMNG; Anatrace) and 0.1% (w/v) cholesteryl hemisuccinate tris salt (CHS; Anatrace) at 4[°C for 1 h. Insoluble components were removed by centrifugation at 70,000g for 30 min. The supernatant was supplemented with 10 mM imidazole and loaded onto a 5 ml Ni-NTA column (GE Healthcare). Beads were first washed with HBS containing 0.01% (w/v) LMNG and 0.001% (w/v) CHS then with HBS containing 0.01% (w/v) LMNG, 0.001% (w/v) CHS and 20 mM imidazole. Proteins were eluted using HBS with 0.01% (w/v) LMNG, 0.001% (w/v) CHS and 250 mM imidazole, and tags were removed using TEV protease. For purification of TauT, the eluate was subjected to SEC using a Superose 6 Increase 10/300 GL column (GE Healthcare) in buffer containing 20 mM HEPES pH 7.4, 150 mM NaCl, 0.01% (w/v) LMNG and 0.001% (w/v) CHS. The eluted peak corresponding to TauT was collected for cryo-EM sample preparation.

### Cryo-EM sample preparation and data acquisition

TauT_APO_, TauT_TAU_ (supplemented with 1 mM taurine), TauT_GABA_ (with 1 mM GABA), TauT_βA_ (supplemented with 1 mM β-alanine) and TauT_GES_(supplemented with 200 μM GES) were diluted to[9 mg/ml and aliquots (3 μl) were applied to GF-0.6/1.0 300 Au-flat (Electron Microscopy Sciences). The cryo-EM grids were prepared using Thermo Fisher Vitrobot Mark IV at 8 °C and 100% humidity (Thermo Fisher Scientific), and data were collected in a Titan Krios microscope (Thermo Fisher Scientific) at 300 kV for data acquisition. Images were collected using a K3 camera (Gatan) mounted post a Quantum energy filter with a 20[eV slit and operated in super-resolution mode with a pixel size of 0.67 Å at the object plane. Defocus values were set to range from −1.5 μm to −2.0 μm for data collection. Data were acquired using EPU-2.9.0.1519REL. The dose rate on the detector was 15.0 e^-^s^-1^A^-2^ with a total exposure of 56 e^-^A^-2^. Each 2.56 s movie was dose-fractioned into 32 frames.

### Image processing

CryoSPARC software was used for structural analysis^46^. The original movies were gain-corrected, motion-corrected and binned to a pixel size of 0.67 Å in a Patch motion correction step. Dose-weighted micrographs were used for CTF estimation using Patch-CTF in cryoSPARC. Particles were auto-picked using Topaz-0.2.3^47^, extracted with a pixel size of 2.68 Å and subjected to two-dimensional (2D) classification to remove contaminants. Particles showing obvious secondary structure features were re-extracted with a pixel size of 1.34 Å and subject to several rounds of ab-initio reconstructions. Seed-facilitated 3D classification was performed to improve the resolution^48^, by several rounds of heterogenous refinement and ab-initio reconstructions. Final good particles were re-extracted with a pixel size of 0.67 Å. Non-uniform and local refinement were performed, and resolutions were estimated by gold-standard Fourier shell correlation.

### Model building

Alphafold^49^ was used to build initial models of TauT and docked into the cryo-EM map with UCSF Chimera-1.14^50^. Models were manually rebuilt in Coot-0.9.2^51^ and further refined by Phenix1.19.2-4158^52^. CIF files for ligands were generated in Phenix using eLBOW. The residues contained in the final models are indicated in Extended Data Table 1. Figures were prepared with Pymol-1.7.0.5 (Schrodinger, LLC.) and UCSF ChimeraX-1.5^51^.

### Flow cytometry

293F cells were transfected with HA tagged wild-type and mutant constructs as previously described^53^. Cells were collected 27 h post-transfection and incubated with PE conjugated HA tag antibody (BioLegend, cat# 901518) for 30 min at room temperature, followed by analysis on a flow cytometer (BD LSRFortessa^TM^). FACS data were analyzed using FlowJo v10.6.2 software.

### Quantification and statistical analysis

Global resolution estimations of cryo-EM density maps are based on the 0.143 Fourier Shell Correlation criterion^54^. The local resolution map was calculated using cryoSPARC-4.6.1^46^. Prism GraphPad software v9.3.0 was used for statistical analysis. The number of biological replicates (N) and the relevant statistical parameters for each set of experiments (such as mean or standard error) are described in the figure legends. P values were calculated by two-sided unpaired t-tests. No statistical methods were used to pre-determine sample sizes.

## Extended data figures and tables

**Extended Data Fig 1.** Sequence alignment and purification of hTauT. **a**, Sequence alignment of Homo sapiens TauT, Mus musculus TauT, Bos taurus TauT, Homo sapiens SLC6A1 and Homo sapiens SLC6A8. Secondary structures of TauT are shown above the alignment with TM1-5, TM6-10, TM11-12, extracellular helices and cytosolic helices colored in blue, green, orange, yellow and pink, respectively. Unmodeled residues are shown as dashed lines. Non-conserved residues are highlighted in gray. Alignment was made using PROMALS3D. **b**, Time-dependent accumulation of [^3^H]taurine in oocytes. Accumulation of oocytes injected with (blue) and without (red) cRNA of wild-type human TauT. Data are shown as mean ± SD (n=3 biologically independent experiments). **c**, The dose-dependent inhibition of hTauT by taurine. The IC_50_ value for taurine of wild-type hTauT is 2.63 ±1.04 µM. Data are shown as mean ± SD (n=3 biologically independent experiments). **d**, Size-exclusion chromatography profile and representative SDS-PAGE analysis of hTauT in detergent. The asterisk indicates the fractions used for cryo-EM sample preparation.

**Extended Data Fig 2.** Data processing of different TauT datasets **a**, Representative raw electron micrograph of TauT. Scale bar, 50 nm. **b**, Two-dimensional class averages of hTauT output from cryoSPARC. **c-d**, Flow chart showing data processing of hTauT_APO_ and ligand-bound hTauT.

**Extended Data Fig 3.** Cryo-EM analysis of TauT in different states. **a-e**, Resolution estimation based on the criterion of the FSC 0.143 cut-off, angular distribution of the final reconstruction and local resolution distribution of hTauT_APO_(a), hTauT_TAU_ (b), hTauT_βA_ (c), hTauT_GABA_ (d) and hTauT_GES_ (e), respectively.

**Extended Data Fig 4.** Cryo-EM density of hTauT in the different states. **a**, Cryo-EM density corresponding to the TM1-12, N-terminal and C-terminal of TauT_TAU_. **b-c**, Cryo-EM density corresponding to the TM1 of TauT_APO_ and TauT_GES_.

**Extended Data Fig 5.** Structural details in the inward-facing conformation. **a,** Cut-open view of electrostatic surface of hTauT_APO_ shows the substrate translocation pathway. **b,** Close-up view of the residues in extracellular gate. Hydrogen bonds and salt bridges are indicated by dashed lines. **c,** Cryo-EM density corresponding to conserved chloride binding site. Residues are shown as sticks. Cl^−^ ion is shown as dot. Coordination of Cl^−^ is indicated by dashed lines.

**Extended Data Fig 6.** Structural analysis of the Na1, Na2 and chloride binding sites. **a-c,** Cryo-EM density corresponding to Na1 binding site of hTauT_TAU_ (a), hTauT_βA_ (b) and hTauT_GABA_ (c). Residues and ligands are shown as sticks. Na^+^ ion is shown as dot. **d-f,** Cryo-EM density corresponding to Cl^−^ binding site of hTauT_TAU_ (d), hTauT_βA_ (e) and hTauT_GABA_ (f). Residues and ligands are shown as sticks. Cl^−^ ion is shown as dot. **g-i,** Cryo-EM density corresponding to Na2 binding site of hTauT_TAU_ (g), hTauT_βA_ (h) and hTauT_GABA_ (i). Residues and ligands are shown as sticks. Taurine-like and β-Alanine-like densities are colored in yellow. Na^+^ densities are colored in purple. **j,** Superposition of Na1 binding site of hTauT_APO_ with hTauT_TAU_. Na^+^ ion is shown as dot and surrounding residues are shown as sticks. Shift of Cα of F58 is indicated by the arrow. hTauT_APO_ and hTauT_TAU_ are colored in light peach and pink, respectively. **k,** Superposition of Na2 binding site of hTauT_APO_ with hTauT_GABA_. Na^+^ ion is shown as dot and surrounding residues are shown as sticks. Shift of Cα of G56 and V59 are indicated by the arrow. hTauT_APO_ and hTauT_GABA_ are colored in light peach and orange, respectively. **l,** Superposition of Cl binding site of hTauT_APO_ with hTauT_TAU_. Cl^−^ ion is shown as dot and surrounding residues are shown as sticks. hTauT_APO_ and hTauT_TAU_ are colored in light peach and pink, respectively.

**Extended Data Fig 7.** Structure comparison of TauT with related transporters. **a,** Superposition of hTauT_GABA_ (orange) with GABA-bound GAT1 (grey). Close-up view shows the binding pocket of GABA. GABA and surrounding residues are shown as sticks. Na^+^ and water molecules are shown as dot. **b,** Superposition of hTauT_GABA_ (orange) with nipecotic acid-bound GAT1 (grey). Close-up view shows the binding pocket of GABA and nipecotic acid. GABA, nipecotic acid and surrounding residues are shown as sticks. Na^+^ and water molecules are shown as dot. **c,** Superposition of hTauT_GES_ (green) with tiagabined-bound GAT1 (grey). Close-up view shows the binding pocket of GES and tiagabine. GES, tiagabine and surrounding residues are shown as sticks.

