## Supplemental figure for "Structure of the human taurine transporter TauT reveals substrate recognition and mechanisms of inhibition"

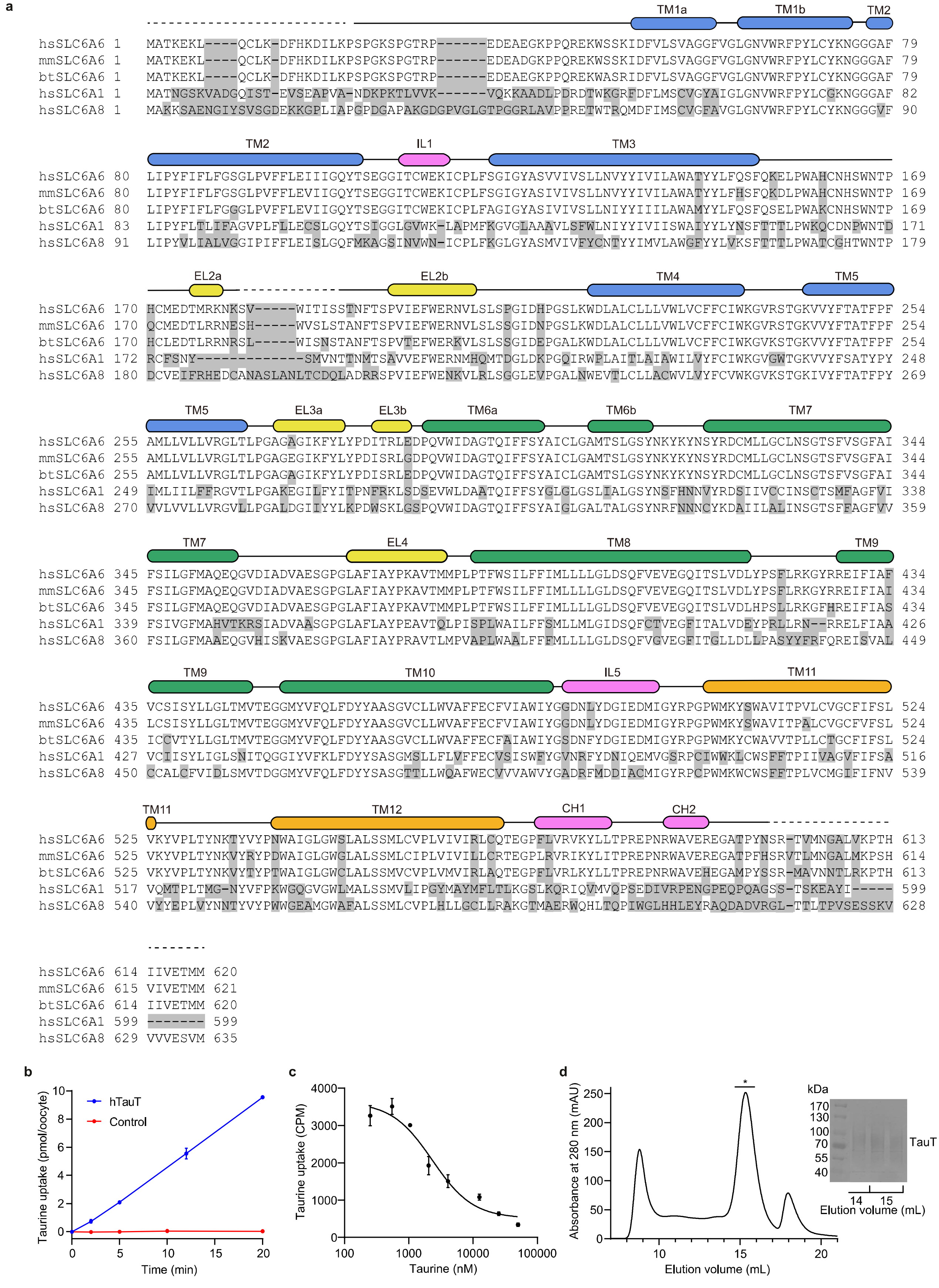


**Extended Data Fig 1. Sequence alignment and purification of hTauT.**

**a,** Sequence alignment of Homo sapiens TauT, Mus musculus TauT, Bos taurus TauT, Homo sapiens SLC6A1 and Homo sapiens SLC6A8. Secondary structures of TauT are shown above the alignment with TM1-5, TM6-10, TM11-12, extracellular helices and cytosolic helices colored in blue, green, orange, yellow and pink, respectively. Unmodeled residues are shown as dashed lines. Non-conserved residues are highlighted in gray. Alignment was made using PROMALS3D.

**b,** Time-dependent accumulation of [$^3$H]taurine in oocytes. Accumulation of oocytes injected with (blue) and without (red) cRNA of wild-type human TauT. Data are shown as mean ± SD (n=3 biologically independent experiments).

**c,** The dose-dependent inhibition of hTauT by taurine. The $IC_{50}$ value for taurine of wild-type hTauT is 2.63 ±1.04 µM. Data are shown as mean ± SD (n=3 biologically independent experiments).

**d,** Size-exclusion chromatography profile and representative SDS-PAGE analysis of hTauT in detergent. The asterisk indicates the fractions used for cryo-EM sample preparation.

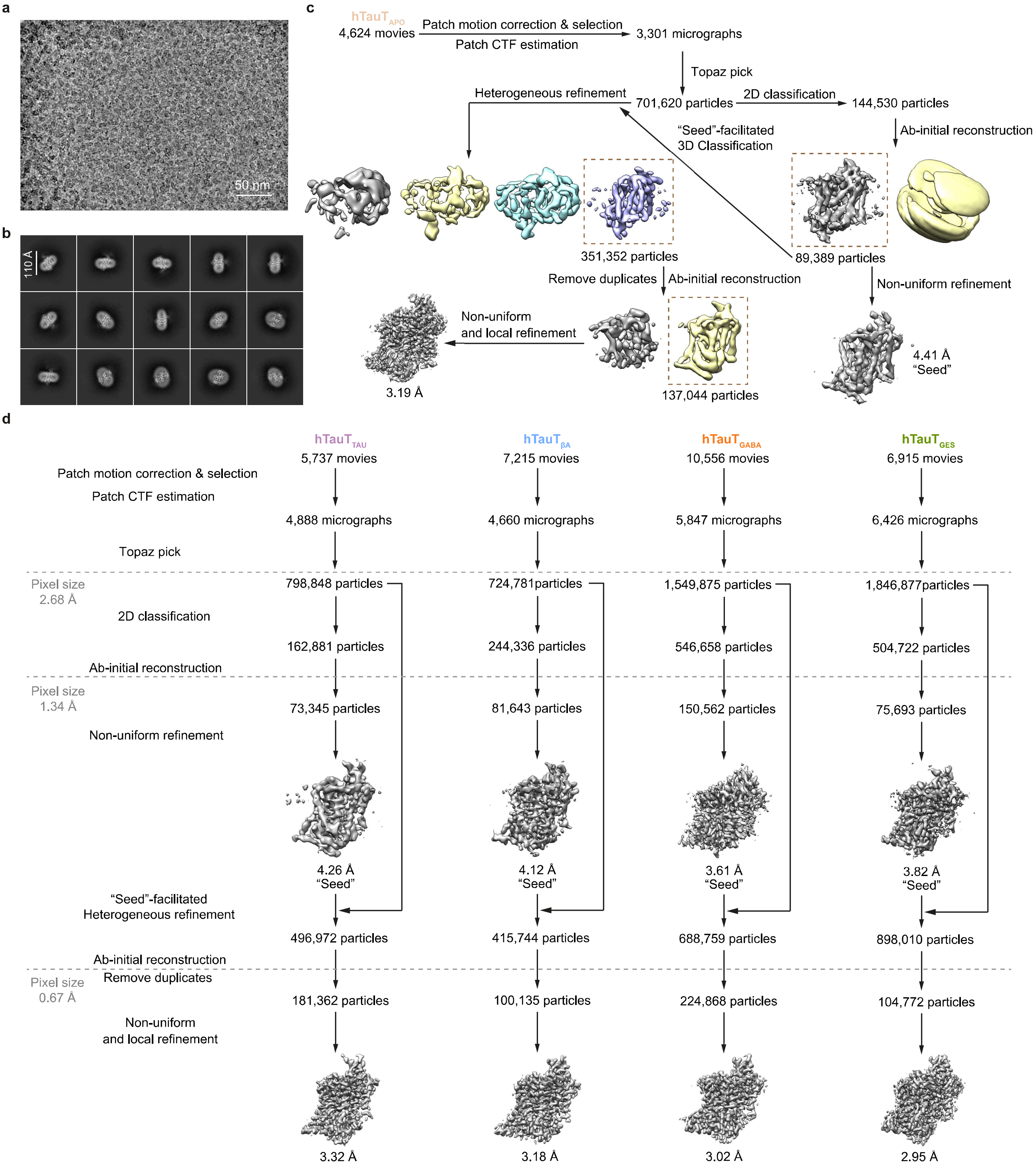


**Extended Data Fig 2. Data processing of different TauT datasets**

**a,** Representative raw electron micrograph of TauT. Scale bar, 50 nm.

**b,** Two-dimensional class averages of hTauT output from cryoSPARC.

**c-d,** Flow chart showing data processing of hTauTAPO (c) and ligand-bound hTauT datasets (d).

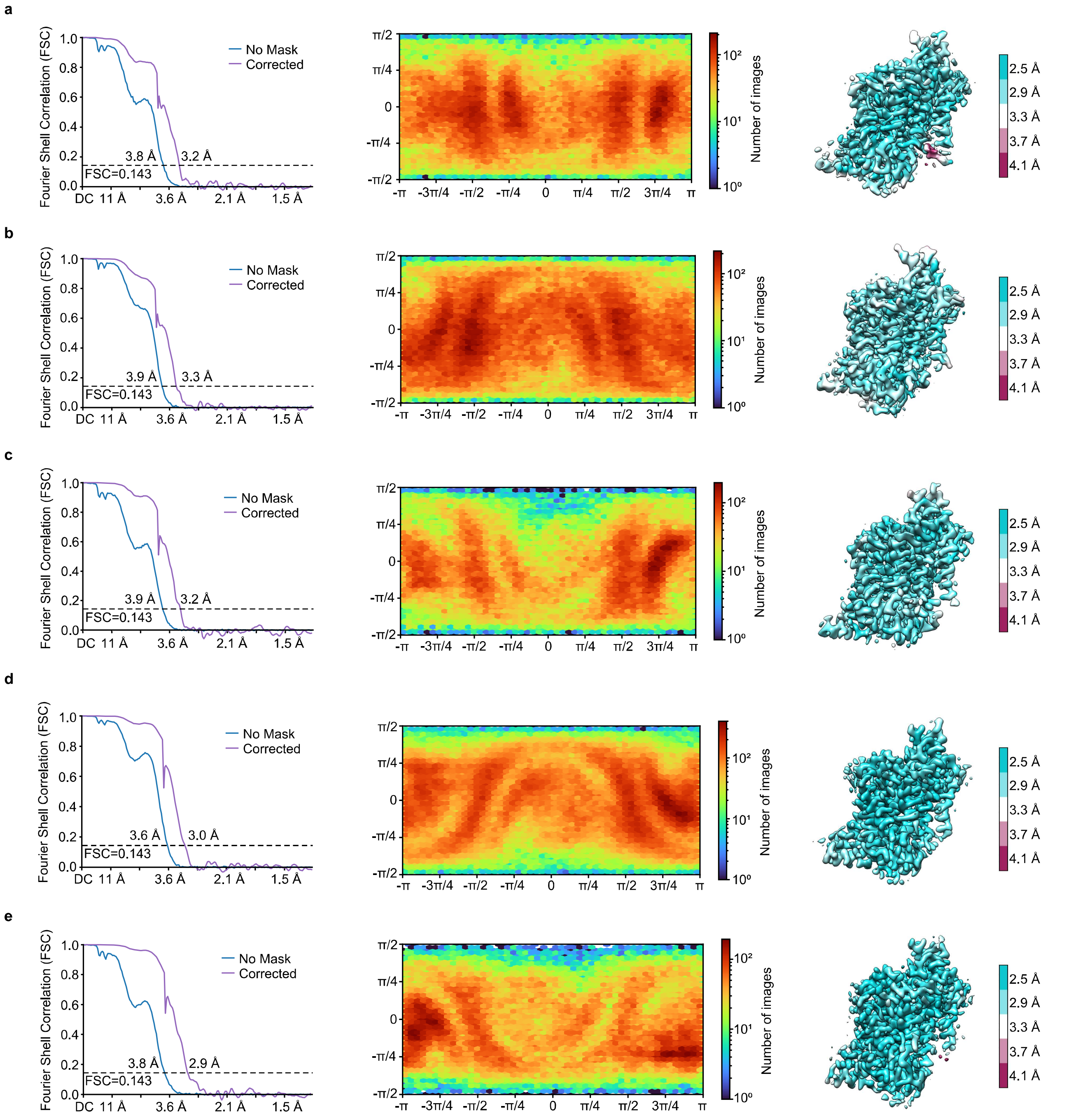


**Extended Data Fig 3. Cryo-EM analysis of TauT in different states.**

**a-e,** Resolution estimation based on the criterion of the FSC 0.143 cut-off, angular distribution of the final reconstruction and local resolution distribution of $hTauT_{APO}$ (a), $hTauT_{TAU}$ (b), $hTauT_{\beta A}$ (c), $hTauT_{GABA}$ (d) and $hTauT_{GES}$ (e), respectively.

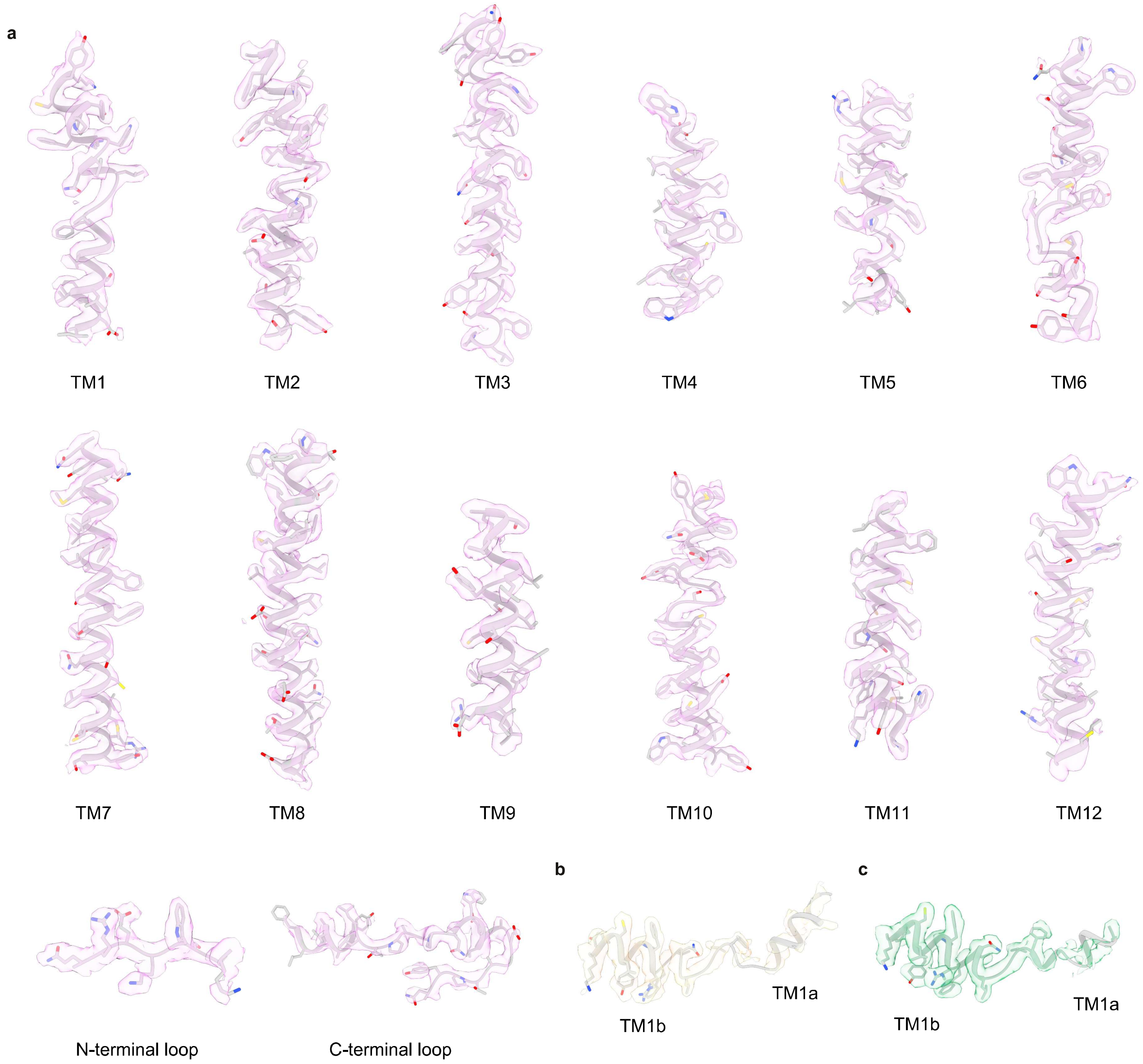


**Extended Data Fig 4. Cryo-EM density of hTauT in the different states.**
**a,** Cryo-EM density corresponding to the TM1-12, N-terminal and C-terminal of $hTauT_{TAU}$.
**b-c,** Cryo-EM density corresponding to the TM1 of $hTauT_{APO}$ and $hTauT_{GES}$.

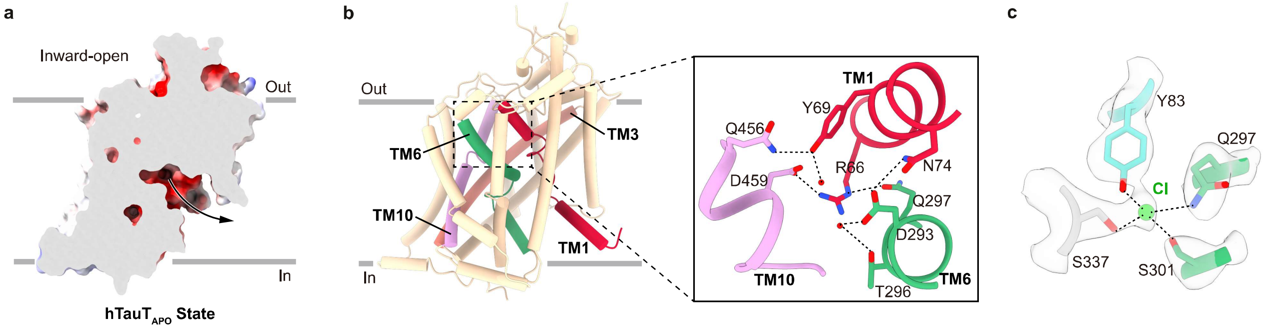


**Extended Data Fig 5. Structural details in the inward-facing conformation.**
**a,** Cut-open view of electrostatic surface of $hTauT_{APO}$ shows the substrate translocation pathway.
**b,** Close-up view of the residues in extracellular gate. Hydrogen bonds and salt bridges are indicated by dashed lines.
**c,** Cryo-EM density corresponding to conserved chloride binding site. Residues are shown as sticks. $Cl^{-}$ ion is shown as dot. Coordination of $Cl^{-}$ is indicated by dashed lines.

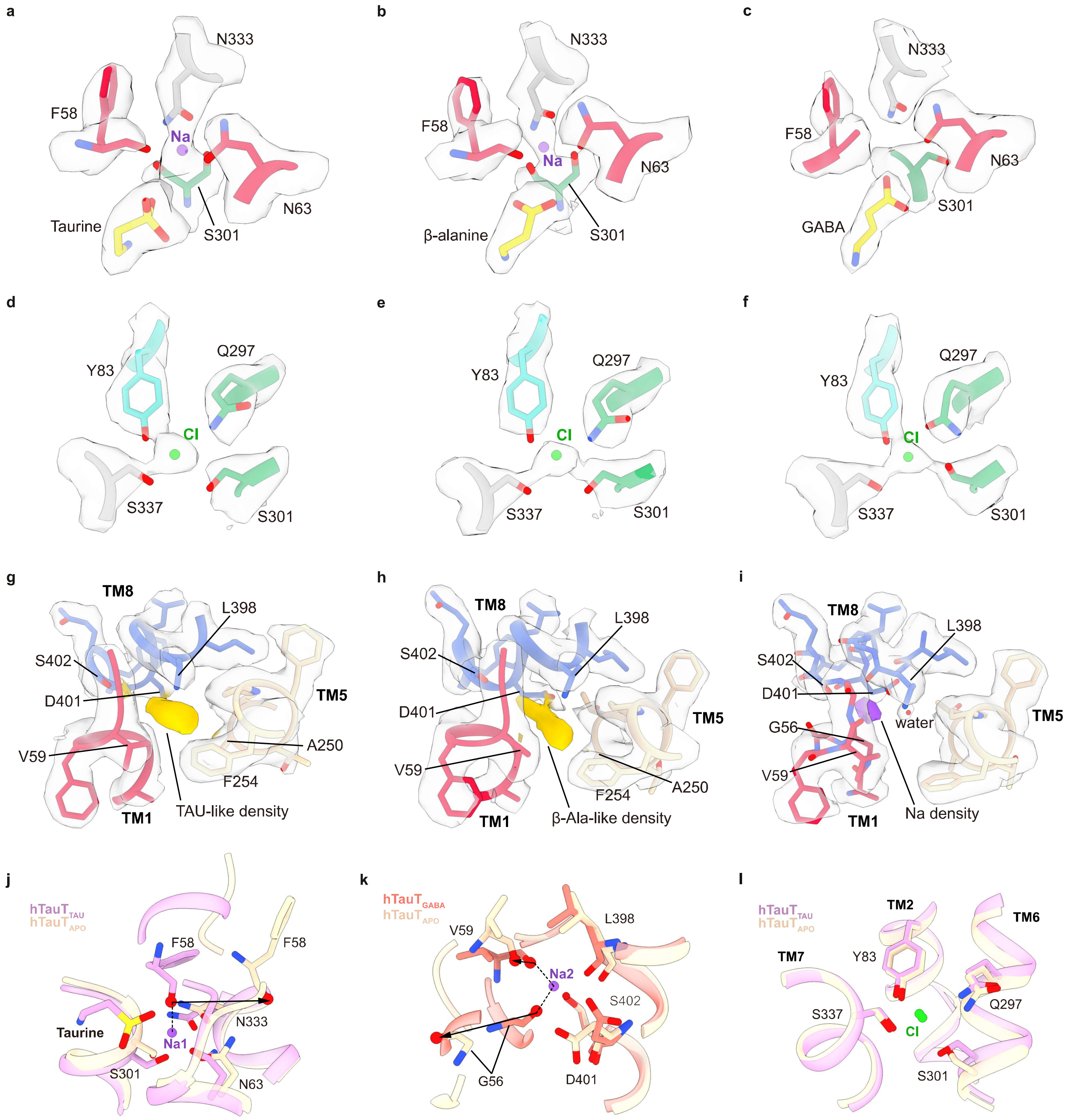


**Extended Data Fig 6. Structural analysis of the ion binding sites in different states.**

**a-c,** Cryo-EM density corresponding to Na1 binding site of $hTauT_{TAU}$ (a), $hTauT_{βA}$ (b) and $hTauT_{GABA}$ (c). Residues and ligands are shown as sticks. $Na^+$ ion is shown as dot.

**d-f,** Cryo-EM density corresponding to $Cl^-$ binding site of $hTauT_{TAU}$ (d), $hTauT_{βA}$ (e) and $hTauT_{GABA}$ (f). Residues and ligands are shown as sticks. $Cl^-$ ion is shown as dot.

**g-i,** Cryo-EM density corresponding to Na2 binding site of $hTauT_{TAU}$ (g), $hTauT_{βA}$ (h) and $hTauT_{GABA}$ (i). Residues and ligands are shown as sticks. Taurine-like and β-Alanine-like densities are colored in yellow. $Na^+$ density is colored in purple.

**j,** Superposition of Na1 binding site of $hTauT_{APO}$ with $hTauT_{TAU}$. $Na^+$ ion is shown as dot and surrounding residues are shown as sticks. Shift of Cα of F58 is indicated by the arrow. $hTauT_{APO}$ and $hTauT_{TAU}$ are colored in light peach and pink, respectively.

**k,** Superposition of Na2 binding site of $hTauT_{APO}$ with $hTauT_{GABA}$. $Na^+$ ion is shown as dot and surrounding residues are shown as sticks. Shift of Cα of G56 and V59 are indicated by the arrow. $hTauT_{APO}$ and $hTauT_{GABA}$ are colored in light peach and orange, respectively.

**l,** Superposition of Cl binding site of $hTauT_{APO}$ with $hTauT_{TAU}$. $Cl^-$ ion is shown as dot and surrounding residues are shown as sticks. $hTauT_{APO}$ and $hTauT_{TAU}$ are colored in light peach and pink, respectively.

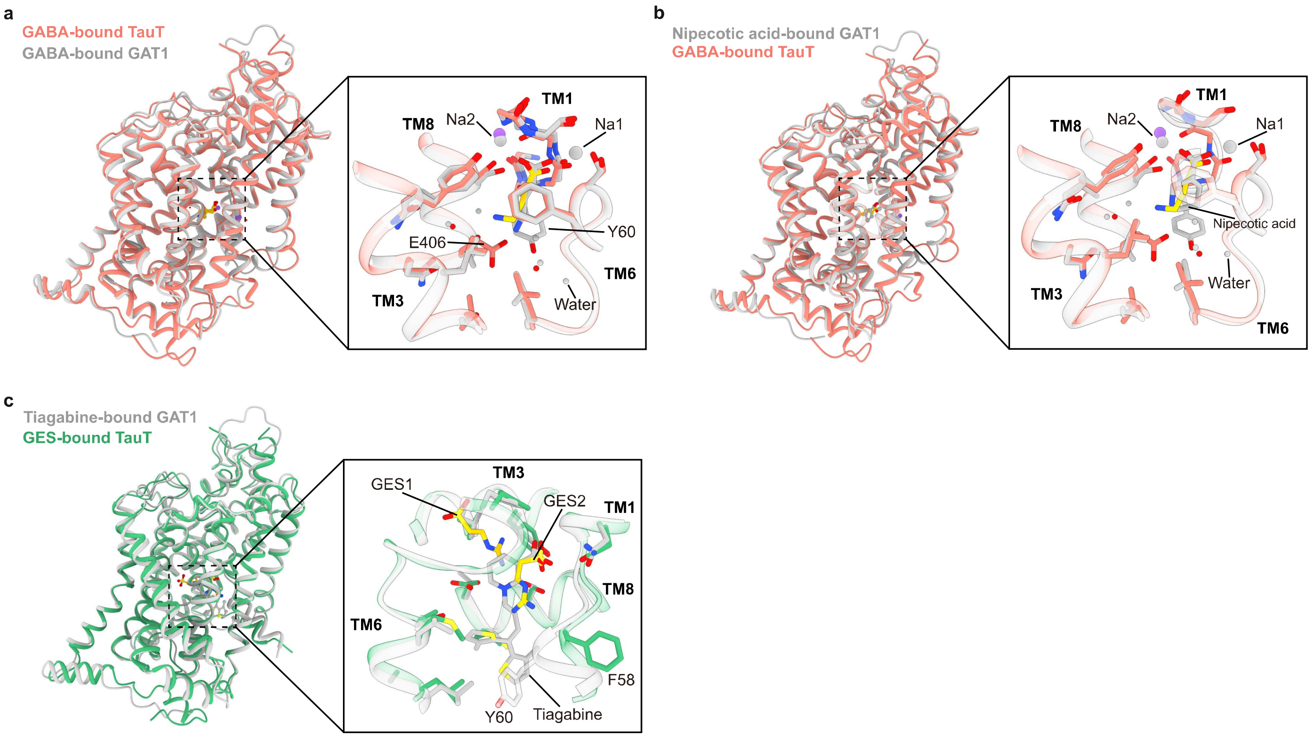


**Extended Data Fig 7. Structure comparison of TauT with related transporters.**
**a,** Superposition of $hTauT_{GABA}$ (orange) with GABA-bound GAT1 (grey). Close-up view shows the binding pocket of GABA. GABA and surrounding residues are shown as sticks. $Na^+$ and water molecules are shown as dot.
**b,** Superposition of $hTauT_{GABA}$ (orange) with nipecotic acid-bound GAT1 (grey). Close-up view shows the binding pocket of GABA and nipecotic acid. GABA, nipecotic acid and surrounding residues are shown as sticks. $Na^+$ and water molecules are shown as dot.
**c,** Superposition of $hTauT_{GES}$ (green) with tiagabined-bound GAT1 (grey). Close-up view shows the binding pocket of GES and tiagabine. GES, tiagabine and surrounding residues are shown as sticks.
