## Supplemental Table 1 for "Structure of the human taurine transporter TauT reveals substrate recognition and mechanisms of inhibition"

**Extended Data Table 1**
**Cryo-EM data collection, refinement and validation statistics**

| | $hTauT_{APO}$ | $hTauT_{TAU}$ | $hTauT_{\beta A}$ | $hTauT_{GABA}$ | $hTauT_{GES}$ |
|---|---|---|---|---|---|
| PDB ID | 9KTS | 9KTT | 9KTU | 9KTV | 9KTX |
| EMDB ID | EMD-62564 | EMD-62565 | EMD-62566 | EMD-62567 | EMD-62569 |
| **Data collection and processing** | | | | | |
| Magnification | 130,000 × | 130,000 × | 105,000 × | 130,000 × | 130,000 × |
| Voltage (kV) | 300 | 300 | 300 | 300 | 300 |
| Electron exposure ($e^-$/Å$^2$) | 56 | 56 | 56 | 56 | 56 |
| Defocus range (μm) | -1.5 to -2.0 | -1.5 to -2.0 | -1.5 to -2.0 | -1.5 to -2.0 | -1.5 to -2.0 |
| Pixel size (Å) | 0.67 | 0.67 | 0.67 | 0.67 | 0.67 |
| Symmetry imposed | *C1* | *C1* | *C1* | *C1* | *C1* |
| Initial particle images (no.) | 701,620 | 798,848 | 724,781 | 1,549,875 | 1,846,877 |
| Final particle images (no.) | 137,044 | 181,362 | 100,135 | 224,868 | 104,772 |
| Map resolution (Å) | 3.2 | 3.3 | 3.2 | 3.0 | 2.9 |
| FSC threshold | 0.143 | 0.143 | 0.143 | 0.143 | 0.143 |
| Map resolution range (Å) | 250-3.2 | 250-3.3 | 250-3.2 | 250-3.0 | 250-2.9 |
| **Refinement** | | | | | |
| Initial model used (PDB code) | P31641-F1 | P31641-F1 | P31641-F1 | P31641-F1 | P31641-F1 |
| Model resolution (Å) | 3.3 | 3.4 | 3.4 | 3.2 | 3.1 |
| FSC threshold | 0.5 | 0.5 | 0.5 | 0.5 | 0.5 |
| Model resolution range (Å) | 250-3.3 | 250-3.4 | 250-3.4 | 250-3.2 | 250-3.1 |
| Map sharpening *B* factor (Å$^2$) | -112.9 | -146.2 | -115.5 | -70.0 | -103.4 |
| Model composition | | | | | |
| Non-hydrogen atoms | 4,228 | 4,335 | 4,212 | 4,358 | 4,310 |
| Protein residues | 537 | 549 | 530 | 548 | 541 |
| Ligands | Cl:1 | Na:1<br>Cl:1<br>TAU:1 | Na:1<br>Cl:1<br>β-Ala:1 | Na:1<br>Cl:1<br>GABA:1 | Cl:1<br>GES:2 |
| *B* factors (Å$^2$) | | | | | |
| Protein | 38.82 | 36.34 | 42.20 | 58.72 | 43.73 |
| Ligand | 34.90 | 25.04 | 42.85 | 61.43 | 46.49 |
| R.m.s. deviations | | | | | |
| Bond lengths (Å) | 0.002 | 0.003 | 0.003 | 0.003 | 0.003 |
| Bond angles (°) | 0.467 | 0.479 | 0.482 | 0.441 | 0.471 |
| Validation | | | | | |
| MolProbity score | 1.37 | 1.72 | 1.68 | 1.64 | 1.44 |
| Clashscore | 3.95 | 6.77 | 5.49 | 4.87 | 4.46 |
| Poor rotamers (%) | 1.12 | 2.19 | 2.00 | 1.95 | 1.33 |
| Ramachandran plot | | | | | |
| Favored (%) | 97.18 | 97.61 | 97.13 | 97.04 | 97.39 |
| Allowed (%) | 2.82 | 2.39 | 2.87 | 2.96 | 2.61 |
| Disallowed (%) | 0.00 | 0.00 | 0.00 | 0.00 | 0.00 |
